# Structural variants contribute to the gut microbiome, blood metabolic traits, and their causal relationships

**DOI:** 10.1101/2025.03.15.643427

**Authors:** Qun Zhou, Keqing Wang, Junhong Chen, Xinjiang Tan, Xun Xu, Xin Jin, Shaoqiao Zhang, Liang Xiao, Tao Zhang, Xiaomin Liu

## Abstract

Research on genetic variants affecting the gut microbiome and blood metabolites has mainly focused on single nucleotide variants (SNPs), with structural variants (SVs) remaining largely unexamined. In this study, we utilized an advanced structural variant detection pipeline on whole-genome sequencing data from 2,002 individuals with an average sequencing depth of 42 folds, identifying 138,859 high-confidence SVs, 49.57% of which were previously unreported. We performed genome-wide association studies on 22,519 common SVs with 616 gut microbial features and 121 blood metabolic traits, uncovering 30 significant links between SVs and microbial taxa/pathways and 48 connections between SVs and blood metabolites. Notable results include 19 associations reaching study-wide significance, such as an Asian-specific 6.5k deletion at the *TENT2* gene, which is strongly associated with the abundance of *Coprobacillus sp. 29_1* (β=0.158, p=2.5×10^−9^), and a deletion covering *HBA1/HBA2/HBM/HBQ118* linked to nine blood metabolic traits, most significantly with mean corpuscular volume (MCV; β=-0.570, p=1.22×10^−91^). Most SVs showed a greater impact than nearby SNPs in their associations with the microbiome/metabolites. Analysis of rare SVs also identified multiple genes associated with the microbiome/metabolites. Additionally, Mendelian randomization using SVs demonstrated causal relationships, including effects from the microbiome to metabolites, such as *Clostridium ramosum*’s impact on serum uric acid and hyperuricemia, and from metabolites to the microbiome, like the influence of mean corpuscular hemoglobin concentration on MF0064: triacylglycerol degradation. This study provides a detailed SV catalog for the Chinese population and highlights the significant role of SVs in the gut microbiome, blood metabolites, and their causal relationships.

**Highlights:**

- Provides a catalog of 138,859 high-confidence SVs and 49.57% of which were novel
- Identify 30 genome-wide significant “SVs—microbial taxa/pathways” and 48 “SVs—blood metabolites” associations, respectively
- An Asian-specific 6.5k deletion at *TENT2* gene associated with the species *Coprobacillus sp. 29_1*
- A deletion spanning *HBA1/HBA2/HBM/HBQ118* associated with 9 blood metabolic traits
- The majority of SVs demonstrated a more pronounced impact than neighboring SNPs in the associations with microbiome/metabolites
- SVs help to reveal the host-microbe causal relationships such as *Clostridium ramosum’*s causal effects on serum uric acid and hyperuricemia

Graphic abstract Based on a multi-omics cohort of 2,002 individuals comprising high-depth whole genome and metagenomic sequencing data, blood metabolites, and detailed questionnaires, we identified 138,859 high-confidence SVs, uncovering 36 significant associations between SVs and microbial taxa/pathways, as well as 48 associations between SVs and blood metabolites. Mendelian randomization using SVs demonstrated causal relationships, such as *Clostridium ramosum*’s causal effect on serum uric acid and hyperuricemia. This study provides a detailed SV catalog for the Chinese population and highlights the significant role of SVs in the gut microbiome, blood metabolites, and their causal relationships.

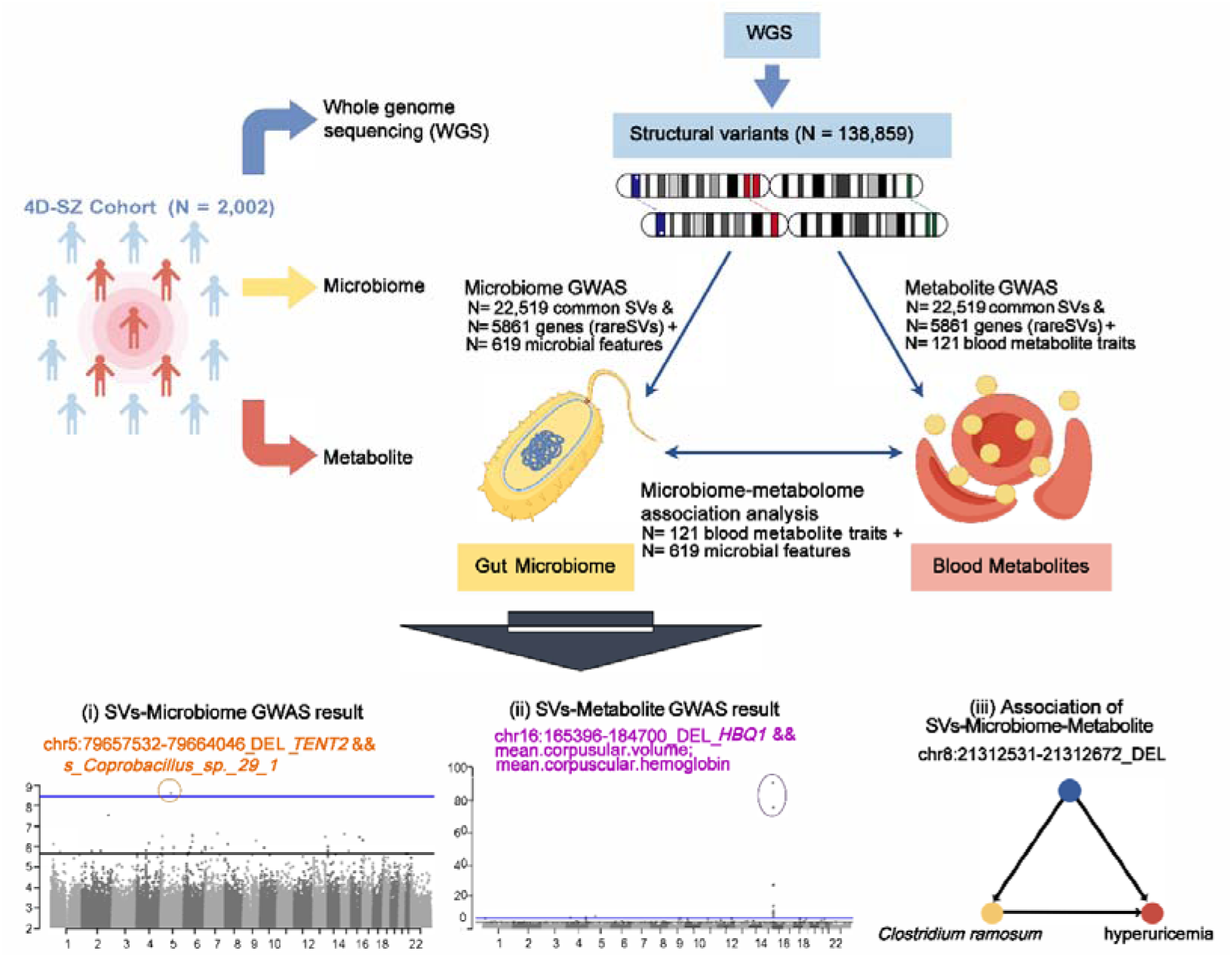

## Introduction

The human gut microbiome plays a pivotal role in numerous physiological processes and is closely linked to human health and the development of diseases such as obesity^1,2^, diabetes^3,4^, inflammatory bowel disease^5,6^, rheumatoid arthritis^7,8^, cardiovascular diseases^9^, and mental disorders^10,11^. Apart from being influenced by factors such as environment, diet, lifestyle, and diseases, the gut microbiome is also controlled to some extent by host genetic factors^12,13^. In recent years, many microbial genome-wide association studies (mbGWAS) have identified various host genetic variants that influence the gut microbiome^14–18^, with only the *LCT* and *ABO* loci being frequently discussed and well replicated.

Despite the progress made in understanding the relationship between host genetics and the gut microbiome, most mbGWAS studies have primarily focused on single nucleotide polymorphisms (SNPs) using genotyping arrays, except for the human microbiome project (HMP) and our prior research^19,20^. This approach has overlooked the potential impact of more extensive and complex genomic variations, such as structural variations (SVs). SVs, which constitute a significant form of genomic variation, including alterations exceeding 50 base pairs (bp) in length such as deletions (DEL), insertions (INS), inversions (INV), duplications (DUP), and copy number variants (CNV). These SVs are prevalent in the human genome and have been associated with various disease phenotypes^21–24^. A recent study identified a correlation between CNV of the human salivary amylase gene (*AMY1*) and the relative abundance of the genus *Ruminococcus* in the gut microbiome^25^. However, this study solely relied on PCR technology to explore the impact of a single human *AMY1* gene copy number variation on the microbiome. Given that few studies had both whole genome and whole metagenome data in the same cohort, thus to date, the impact of host SVs on the gut microbiome remains largely unexplored.

In addition to the gut microbiome, the genome-wide contributions of host SVs to blood metabolic traits have been underexplored. Very limited evidence has been reported. For example, Chen et al. identified a deletion of the *ALB* promoter and a CNV at *PDPR* that were significantly associated with blood metabolic traits in the Finish population^26^. The NHLBI TOPMed program has reported 21 independent SVs or 41 SV-trait associations across a total of 24 quantitative hematologic traits^27^. While these studies have identified associations between SVs and metabolic traits in Finnish and European populations, there is a notable gap in research focusing on Asian populations. Recent efforts to address this gap include a study using long-read sequencing in a Chinese population, which identified 22 SVs on 14 chromosomes associated with 13 phenotypes^28^. However, the association analysis was conducted on a maximum of 327 individuals. This underscores the need for a comprehensive investigation of SV contributions to blood metabolic traits in larger Asian populations. Moreover, comprehensively and systematically excavating the impact of host SVs on both the microbiome and metabolites using high-depth whole-genome data from a well-designed cohort will enhance our understanding of host-microbiome interaction mechanisms.

To address the research gap and investigate the role of SVs in the gut microbiome and blood metabolic traits, we constructed and characterized a high-quality SV profile of the Chinese population based on 2,002 individuals from the 4D-SZ cohort^14,19,20,29–31^, a multi-omics cohort comprising high-depth whole genome and metagenomics sequencing, blood metabolites, and detailed questionnaires. Subsequently, we conducted genome-wide association analyses to identify common SVs and genes harboring rare SVs that significantly contributed to gut microbial taxa/pathways and blood metabolic traits, respectively. Moreover, we examined the intricate relationship between gut microbiome and blood metabolites through their associated SVs as mediators (see **Graphic abstract**). The primary objective of this study was to characterize the landscape of SVs in the Chinese population and identify novel SVs associated with the microbiome and metabolites, while offering valuable insights into the mechanisms governing host-microbiome interactions.

## Results

### Characterization of SVs identified in the Chinese population

We utilized high-depth whole-genome sequencing (WGS) data to identify SVs in 2,002 Chinese individuals from the 4D-SZ cohort. Among the sequenced samples, 53% were females, with an average age of 30 years ranging from 21 to 67 years (**Supplementary Fig. 1**). The mean sequencing depth per individual was 42×, amounting to an average of 122.46 Gb of sequenced data (**Supplementary Fig. 2a**). The base mapping rate varied from 96.74% to 100%, with an average of up to 99.89% (**Supplementary Fig. 2b).** An advanced Genome Analysis Toolkit’s Structural Variant (GATK-SV) pipeline^32,33^ was employed to detect SVs (**Supplementary Fig. 3**; Methods). Following rigorous quality control measures for both samples and variants, a total of 1,839 samples were retained. We detected a total of 138,859 high-confidence SVs with quality scores surpassing 250 or marked as “PASS,” while excluding “BND” variant types (**Supplementary Table 1)**. These SVs comprised 73,288 insertions (INSs), 38,782 deletions (DELs), 25,030 duplications (DUPs), 1,153 complex SVs (CPXs), 378 inversions (INVs), and 228 copy number variations (CNVs) (**Fig. 1a**). The most prevalent SV type was INSs, accounting for 52.78% of all SVs, followed by DELs, representing 27.93% (**Fig. 1a**).

**Figure 1.**
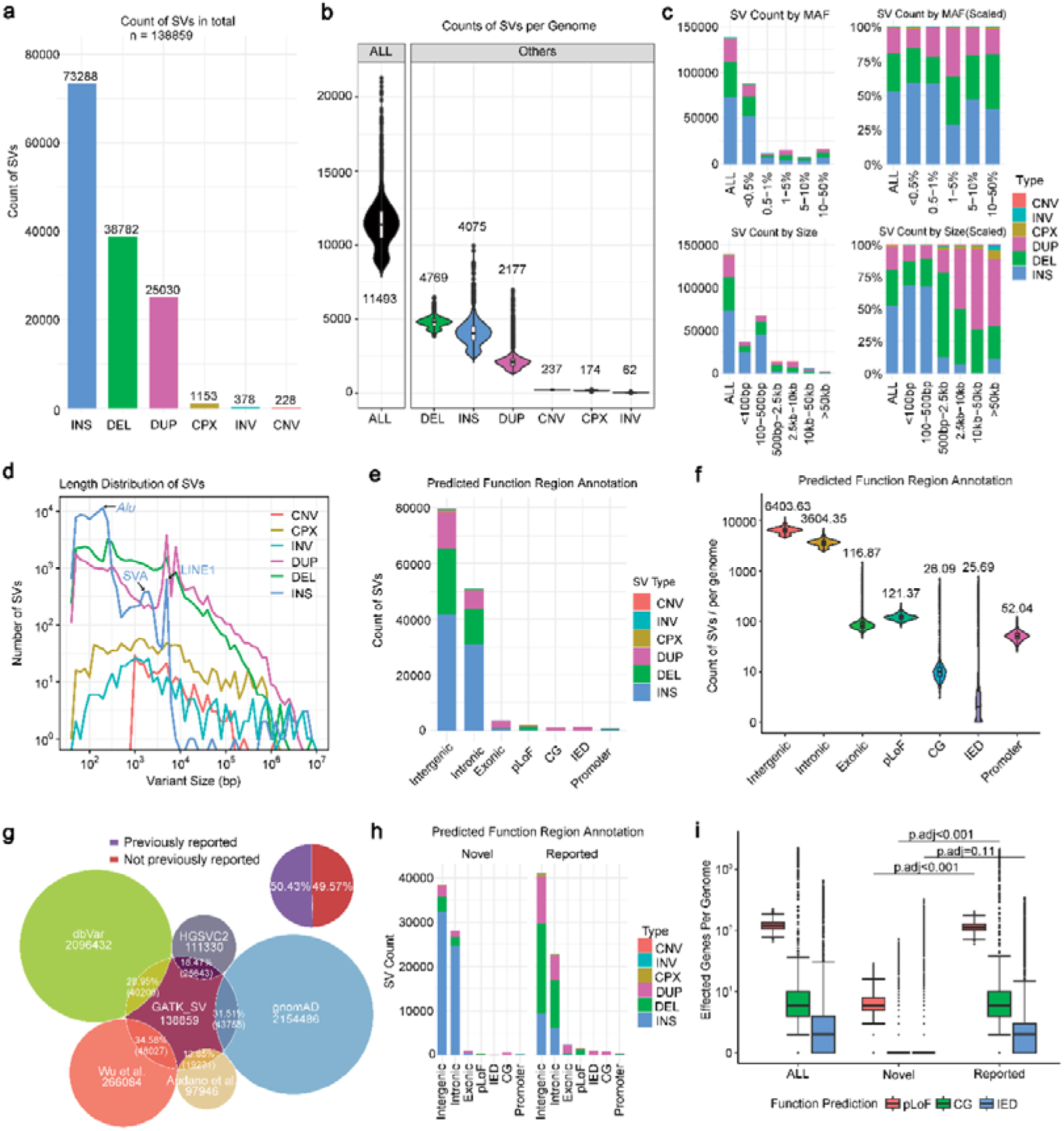
Characteristics of the SVs discovered in this study (4D-SZ Cohort). (a) Descriptive statistics of the SVs count for each SV type. (b) The count of SVs per sample by each SV type. (c) SV count distribution by minor allele frequency (MAF) and size. (d) Length distribution of SVs. The peaks represent the Alu, SVA, and LINE1 events. (e) The count of SVs according to the predicted function region annotation by each SV type. (f) The count of SVs per sample according to the predicted function region annotation by each SV type. (g) Venn diagram showing the overlap between SVs in this study and published SV datasets. Among the SVs, 49.57% are novel, while 50.43% have been previously reported. (h) The difference in SV counts between novel and reported SVs according to the predicted function region annotation. (i) The counts of affected genes per genome by novel and reported SVs according to the predicted function region annotation. Differences between novel and reported SVs were assessed with two-sided t-tests. CPXs were not included in this analysis due to a lack of accurate annotation of the genes they affected. Abbreviations: SV, structural variant; DEL, deletions; DUP, duplications; INS, insertions; INV, inversions; CPX, complex variants; CNV, copy number variation.

Despite the overall higher count of INSs than DELs in the population, each individual exhibited a greater number of DELs than INSs, consistent with an earlier study^34^. On average, each genome harbored 11,493 SVs events (ranging from 7,851 to 21,324), exceeding the SV count documented in the 1000 genome phase3 (9,679 SVs; average depth of 30×) and the Genome Aggregation Database (gnomAD^32^; 7,439 SVs; average depth of 32×), yet lower than a long-read sequencing study^28^ (18,489 SVs; average depth of 17×). DELs and INSs continued to be the most prevalent SV types, constituting 41.5% and 35.5% of all identified SVs, respectively. The average numbers of DELs, INSs, DUPs, CNVs, CPXs, and INVs per sample were 4,769, 4,075, 2,177, 237, 174, and 62, respectively (**Fig. 1b**).

To evaluate the identified SVs, we applied the same sequencing and analysis methods to the cell line obtained from reference sample NA12878 (**Methods**). Through the application of the GATK-SV pipeline, 10,624 SVs were detected. A comparison of the detected SVs with the gold standard truth set of NA12878^35^ revealed a precision of 79.5% for deletions, 64.2% for duplications, 69.4% for insertions, and 75% for inversions. Additionally, the impact of sequencing depth on SV detection was assessed, as shown in the **Supplementary Fig. 4**. Overall, a higher sequencing depth was correlated with an increased number of SVs, with a gradual rise in SV counts from 20× to 50×. However, this trend declined beyond 50×, potentially owing to the limited number of samples used.

Across all population-detected SVs, 71.55% were rare, with minor allele frequencies (MAF) below 1%. Most SVs exhibited lengths distributed within the range of 50∼100bp (61.02%) and 100-500bp (15.77%; **Fig. 1c**). The expected peaks corresponding to the insertion of Alu, SVA, and LINE1 mobile elements were identified at approximately 300 bp, 2 kb, and 6 kb, respectively (**Fig. 1d**). These SV patterns were consistent with those reported in previous studies.

Regarding the functional prediction of SVs, a significant proportion of SVs were located in the intergenic (57.32%) or intronic regions (36.62%; **Fig. 1e**). Only 6.06% of SVs breakpoints intersected with functional elements, including exons (n = 3381), predicted loss-of-function (pLoF; n = 1895), copy gain of entire genes (CG; n=1170), intragenic exonic duplication (IED; n=1308), and promoters (n = 655). On average, each genome harbored a mean of 173.96 SVs that altered the functions of 213 genes (121.37 pLoF affected 122.7 genes, 26.09 CG affected 65.4 genes, and 25.69 IED affected 25.1 genes; **Fig. 1f**), indicating that approximately 1.1% of human protein-coding genes per individual could experience functional abnormalities due to SVs.

To ascertain whether the SVs identified in this study had been previously documented, we compared them with five publicly available SV databases: gnomAD^32^, Audano et al. 2019^36^, the Human Genome Structural Variation Consortium(HGSVC2)^37^, Wu et al.^28^, and dbVar^38^ (accessed on September 30, 2023). Initially, we ensured that the SV coordinates in these databases were converted to GRCh38 to align them with our datasets. An SV in our study was considered previously reported if the overlap with SVs from published datasets was ≥ 50%. Finally, our dataset exhibited the greatest number of overlapping SVs (48,027, overlapping rate 34.58%; **Fig. 1g**) with the dataset constructed by Wu et al., which was based on long-read WGS results from 405 Chinese individuals. The shared ancestry of all Chinese individuals likely contributed to this substantial overlap. Furthermore, a significant number of overlapping SVs were also detected in the gnomAD (43,755; 31.51%) and dbVar (40,206; 28.95%) datasets, two commonly used reference datasets known for their diverse ethnic population representation. In contrast, the overlaps with the HGSVC2 (25,643; 18.47%) and Audano et al. datasets (19,231; 13.85%) were relatively low. Overall, 49.57% (68,830/138,859) of the SVs discovered in our study were novel and had not been previously documented (**Fig. 1g**). These novel SVs were characterized by a higher fraction of insertions (p < 2.2×10^−16^, fisher’s exact test), particularly in intergenic and intronic regions (both p < 2.2×10^−16^, fisher’s exact test; **Fig. 1h)**. Additionally, they showed a lower frequency of overlap with functional elements, affecting an average of 20.61 SV-altered genes per genome, including 6.95 pLoF, 2.24 CG, and 11.41 IED (**Fig. 1i**).

### Associations of SVs count and common SVs with the gut microbiome

After quality control during SV calling, 1,406 of the 1,839 samples had available fecal shotgun metagenomic sequencing data. We first assessed the impact of an individual’s SVs count on gut microbial diversity through SV burden analysis using a generalized linear regression model adjusted for age, gender, and the first 10 principal components. A weak but statistically significant positive correlation was observed between the SV burden and α-diversity indices (Shannon index: β = 0.08, *p* = 0.011; Simpson index: β = 0.06, *p* = 0.012; **Supplementary Fig. 5**). We then calculated the associations of 22,519 common SVs (MAF ≥ 1%) with α-diversity indices and identified no significant associations after multiple test correction (*p* < 2.22 × 10^−6^ = 0.05/22,519), consistent with previous conclusions in the SNP GWAS for α-diversity. The most significant SV was an insertion in the genetic region of chr19:3502006-3502028 associated with the Shanno index (*p*= 7.55 × 10^−5^) and Simpson index (*p*=4.02 × 10^−6^; **Supplementary Fig. 6**). This insertion is near the Fizzy-related protein homolog 1 (*FZR1*) gene, which mainly regulates the cell cycle in association with the anaphase-promoting complex/cyclosome (APC/C), and plays an important role in neurodevelopment^39^ and tumorigenesis^40^.

Next, a microbial genome-wide association study (mbGWAS) was conducted on 22,519 common SVs and 616 microbial features comprising 517 gut microbial taxa and 99 gut metabolic modules (GMMs) present in over 10% of the individuals **(Supplementary Table 2)**. A total of 30 SVs (DEL = 16, INS = 7, DUP = 6, INV=1) were identified to be associated with gut microbial features with genome-wide significance (*p* < 2.22 × 10^−6^; **Fig. 2a; Supplementary Table 3**). The genomic inflation factor (λ) ranged from 0.84 to 1.33, with a median value of 0.98 (**Fig. 2b**), indicating no obvious evidence of inflation, except for some specific mbGWAS tests. Only one association achieved the study-wide significance level (*p* < 3.60 × 10^−9^, adjusting for 22,519 SVs and 616 microbial features).

**Figure 2.**
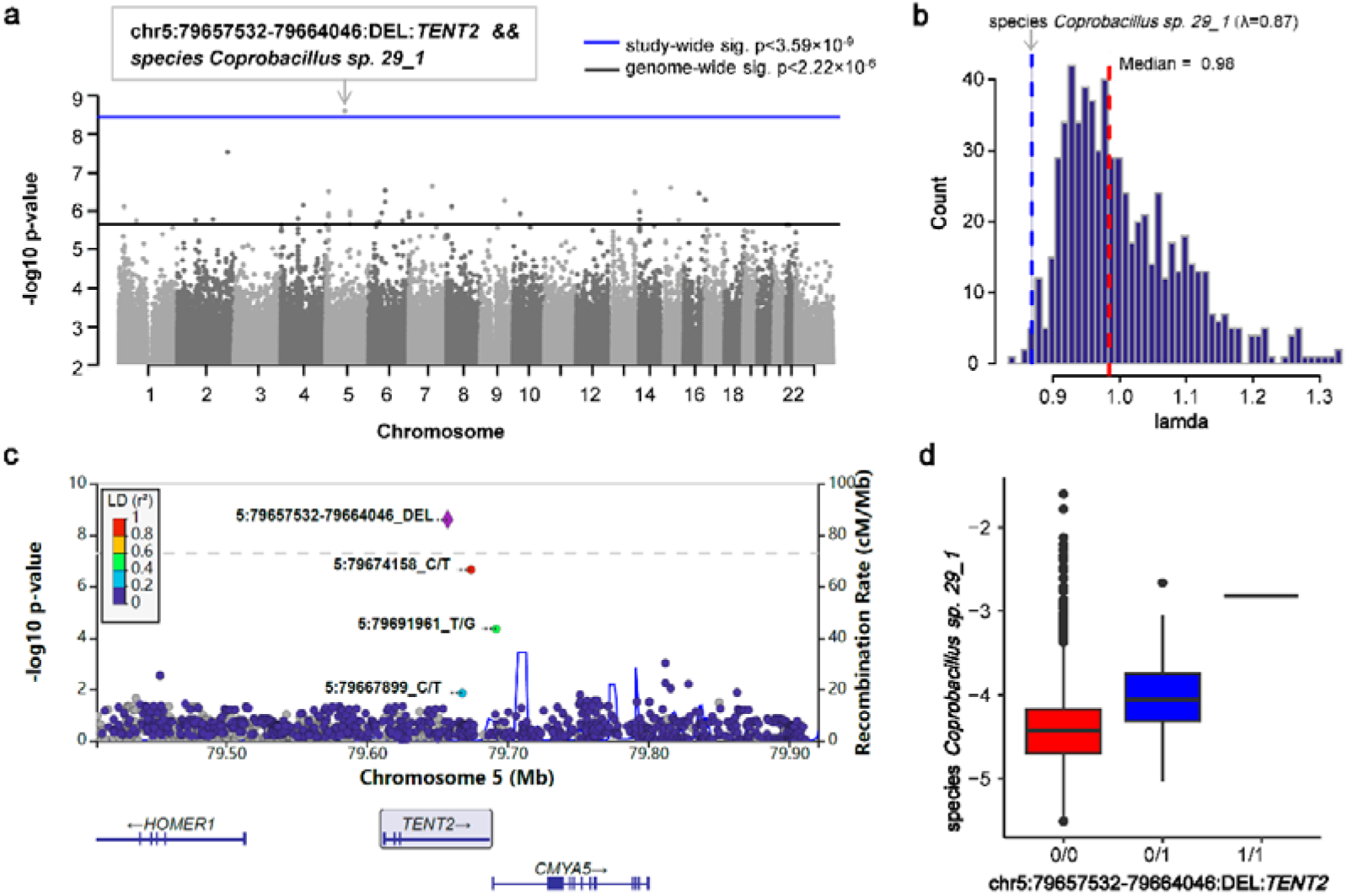
Genome-wide associations of SVs with the gut microbiome identified by mbGWAS. (a) Manhattan plot representing the GWAS results (619 microbial features; 22,519 common high-quality SVs with MAF ≥ 1%). The y-axis indicates the log10-transformed p-values, while the x-axis indicates the genomic positions of the SVs. The study-wide significance (p < 3.59 × 10^−9^) and genome-wide significance (p < 2.2 × 10^−6^) are indicated by horizontal lines colored blue and black, respectively. The annotations highlight the study-wide association of deletion variant at the intronic region of *TENT2* gene and the species *Coprobacillus sp. 29_1*. (b) The distribution of genomic inflation factor (λ) for 616 mbGWAS tests, ranging from 0.84 to 1.33 with a median value of 0.98. The λ of GWAS for species *Coprobacillus sp. 29_1* is 0.87. (c) The regional plot showing mbGWAS results of the *TENT2* deletion (purple diamond) and its nearby SNPs associated with the species *Coprobacillus sp. 29_1*. (d) The log-transformed relative abundances of *Coprobacillus sp. 29_1* in carriers of *TENT*2 deletion and non-carriers.

The most significant association was observed for a 6.5kb deletion (chr5:79657532-79664046) within the intron region of the gene *TENT2*, which was positively correlated with the relative abundance of the species *Coprobacillus sp. 29_1* (β = 0.158, *p* = 2.50 × 10^−9^; **Fig. 2c**). Carriers of this SV exhibited significantly higher mean relative abundances of *Coprobacillus sp. 29_1* (homozygous DEL: 0.15%; heterozygous DEL: 0.024%) than the non-carriers (0.015%; **Fig. 2d**). This GWAS test exhibited no inflation, with a λ value of 0.87. The identified SV had an MAF of 0.014 in this cohort and was specific to East Asian (MAF=0.028 for ASN; MAF=0 for other populations) according to the 1000 Genomes database^41^. Further analysis revealed that this SV was correlated with concentrations of glycine (*p* = 0.015), ethanol (*p* = 0.022), monocytes (*p* = 0.027), and albumin (*p* = 0.028) by investigating its associations with blood parameters in this cohort (**Supplementary Fig. 7**). Interestingly, *Coprobacillus sp. 29_1* associated with SV also showed a correlation with monocyte concentration, suggesting potential interactions among the three. By calculating the pairwise linkage disequilibrium (LD) between this SV and its neighboring SNPs, we found that this SV was in a strong LD with the SNP chr5:79674158_C/T (r^2^ = 0.973; **Fig. 2c**), which also resided in the intronic region of *TENT2* and showed a significant association with *Coprobacillus sp. 29_1* (β = 0.143, *p* = 2.18 × 10^−7^). Consistent with SV, this SNP had a MAF of 0.013 in this cohort and 0.027 in the ASN population, but was absent in other populations from the 1000 Genomes database. This SNP was identified as a susceptibility mutation for schizophrenia^42^, and *Coprobacillus sp. 29_1* has been consistently reported to be enriched in schizophrenia patients^11^. Due to the strong LD between the SV and the SNP, the association with *Coprobacillus sp. 29_1* became insignificant after conditioning on either of them. Both the SV and SNP were located in the intronic region of gene *TENT2*, which is well known to regulate miRNA stability and abundance via diverse mechanisms^43^. The existing literature indicated the significant role of host miRNAs in modulating the gut microbiota^44^. We hypothesized that SV or SNP may affect the gene expression of *TENT2*, potentially regulating miRNA to influence the gut microbial abundance.

In addition to the study-wide significant signal, 22 SVs were associated with microbial taxa at genome-wide significance (**Supplementary Table 3**). Among them, three SVs lied in the genes that were extremely intolerant to pLoF variants with pLI ≥ 0.99, as annotated by the AnnotSV tool. Specifically, a 61 bp duplication within the *CDH8* gene was negatively associated with the abundance of *Streptococcus sanguinis* (β = –0.137, *p* = 3.37 × 10^−7^), a predominant bacterium in the human oral cavity known to be linked to the development of infective endocarditis^45^. Similarly, a 108 bp duplication in the *ANKRD11* gene was negatively associated with the abundance of *Turicibacter* (β = –0.133, *p* = 4.91 × 10^−7^). The SV was overlapped with pathogenic loss of *ANKRD11*, which caused KBG syndrome, a disease characterized by intellectual disability, skeletal malformations, and macrodontia^46^. *Turicibacter* has been reported to be a highly heritable taxon in both human and mouse genetic studies^12,47,48^, supporting the association between SV and *Turicibacter* abundance found in this study. Conversely, a 751 bp deletion in the *GRID2* gene was positively associated with the presence of *Eubacterium cellulosolvens* (β = 0.755, *p* = 6.85 × 10^−7^), an anaerobic cellulolytic bacterium in the gut. The *GRID2* gene encodes the glutamate receptor channel delta-2 subunit, and its deletion leads to a recessive syndrome of cerebellar ataxia and tonic upgaze in humans^49^.

In addition to the three SVs within disease-associated genes, several SVs were found to exhibit significant effects on potential probiotics that could contribute to host health. For example, a duplication event in the *LOC107985978* gene on chromosome 7 (SV length: 68 bp) was positively associated with the abundance of *Roseburia intestinalis* (β = 0.139, *p* = 2.20 × 10^−7^), a promising probiotic known for its anti-inflammatory properties by producing butyrate^50,51^. An insertion on chromosome 15 involving an ALU element (SV length: 262 bp) demonstrated a significantly negative correlation with the emerging probiotic *Parabacteroides distasonis*^52^ (β = –0.137, *p* = 2.51 × 10^−7^).

In addition to microbial taxa, seven associations were identified for the GMMs, also known as microbial pathways. For instance, the glyoxylate bypass pathway (MF0063) was negatively associated with an insertion on chromosome 2 (SV length: 111 bp, β = –0.147, *p* = 2.92 × 10^−8^). The pyruvate dehydrogenase complex (MF0072) exhibited a negative association with a deletion on chromosome 13 (SV length: 234 bp, β = –0.132, *p* = 3.26 × 10^−7^). The pectin degradation II pathway (MF0004) exhibited a negative association with a deletion on chromosome 9 (SV length: 82 bp, β = –0.131, *p* = 5.19× 10^−7^). Notably, all the SVs associated with the microbiome were located in the intronic and intergenic noncoding regions, except for an inversion of 4.78Mb spanning 14 genes, consistent with previous findings in the SNP-based mbGWAS^53^.

### Conditional analyses and functional annotations of microbiome-associated SVs

To further investigate the independence of the identified microbiome-associated SVs from nearby SNPs in influencing changes in microbial features, pairwise LD between SVs and their neighboring SNPs within the ±500kbp flanking region of SVs were calculated. We found that eight of the 30 microbiome-associated SVs exhibited at least moderate LD (r^2^ ≥ 0.5) with one SNP or indel associated with the same microbial feature (**Fig. 3a**; **Supplementary Table 3-4**). Subsequently, conditional analyses were performed for each SV, adjusting for the tagging SNP that showed the strongest LD with SV. These analyses confirmed that eight SV-microbiome associations were no longer significant after adjusting for SNP/indel at the same locus (*p* > 0.01; **Fig. 3b**), whereas the remaining 22 SVs maintained their significance. Notably, 11 of these 22 SVs retained genome-wide significance following conditional analyses (*p* < 2.2×10^−6^). These independent SV signals included two SVs that were potentially linked to the probiotics—*Roseburia intestinalis* and *Parabacteroides distasonis*. These findings indicate that the majority of SVs (73%), rather than nearby SNPs, were the primary causal agents driving changes in microbial abundance.

**Figure 3.**
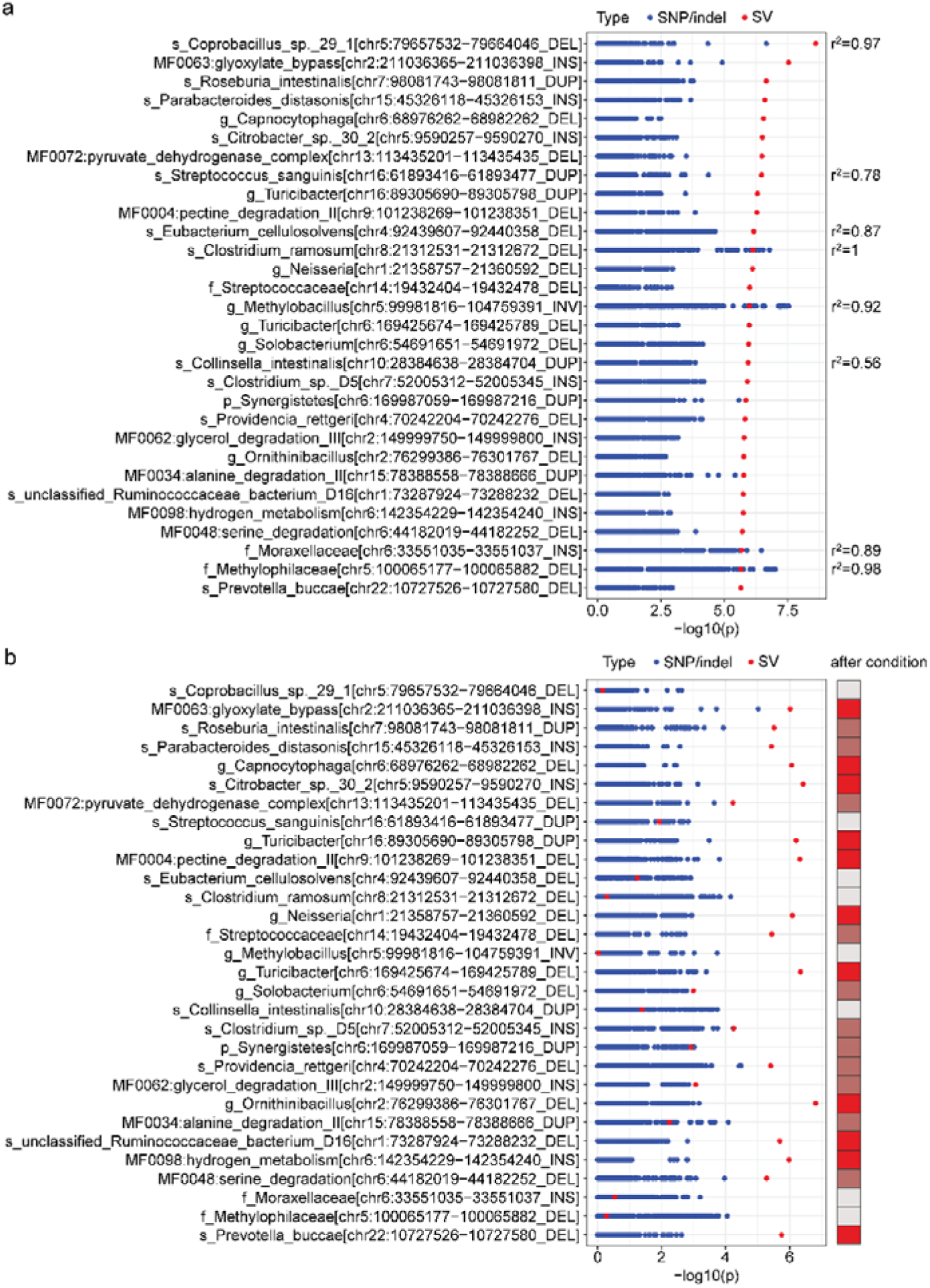
The raw and conditional GWAS results for significant SVs and their nearby SNPs/indels with the gut microbiome. **(a)** The raw associations of SVs and their nearby SNPs/indels (flanking 500kb of the SV) with gut microbiome. **(b)** The associations of SVs and their nearby SNPs after conditioning on the tagging SNP/indel showing the strongest LD with the SV. The bar in the right indicates the significance of SV association after conditional analysis, including no longer significant (colored in grey), not reaching genome-wide significant (colored in dark red) and retaining genome-wide significant (colored in red). The x axis indicates the log10-transformed p values. The y axis indicates the associations of the SVs and nearby SNPs/indels with gut microbiome (N = 30). The red dots represent the p-values associated with SVs, and the blue dots represent the p-values associated with SNPs/indels.

AnnotSV^54–56^ was employed to annotate the 30 microbiome-associated SVs, resulting in the identification of 24 genes that harbored these SVs. Subsequently, gene enrichment analysis was conducted for these 24 genes. Gene Ontology (GO) enrichment analysis revealed 10 significantly enriched GO terms (*p*.adjust < 0.05), including one biological process (BP), six cellular components (CC), and one molecular function (MF) (**Supplementary Fig. 8a**; **Supplementary Table 5**). The enriched gene functions were predominantly related to the structure and function of neuronal synapses, encompassing the synaptic membrane, synaptic cleft, sodium-independent organic anion transport, sodium-independent organic anion transmembrane transporter activity, postsynaptic specialization membrane, postsynaptic density membrane, glutamatergic synapse, and excitatory synapse. Additionally, Kyoto Encyclopedia of Genes and Genomes (KEGG) enrichment analysis indicated significant enrichment of these genes in four pathways: nicotinate and nicotinamide metabolism, long-term depression, the peroxisome, and the phosphatidylinositol signaling system (**Supplementary Fig. 8b**; **Supplementary Table 6**). These findings suggested a pivotal role of microbiome-associated SVs in neurotransmission, ion transport, and cell signaling pathways.

### Associations of rare SVs and the gut microbiome

To further explore the relationship between rare SVs (MAF < 0.01) and the gut microbiome, we performed a gene-based sequence kernel association test (SKAT) analysis of overall diversity and microbial abundances. With a p-value threshold of 8×10^−6^ by adjusting for the detected 5861 genes, we did not observe a significant association between overall diversity and rare SV burden. However, 56 gene loci were associated with microbial features (**Supplementary Table 7**). Two associations reached the study-wide significance (*p*<1.38× 10^−9^ by adjusting for 5861 genes and 616 microbial features). The strongest signal was observed for the Janus kinase and microtubule interacting protein 2 (*JAKMIP2*) gene (harboring four rare SVs) associated with the abundance of MF0103: mucin degradation (*p*=8.28×10^−10^). The *JAKMIP2* gene has been reported as a potential risk factor for Graves’ disease in the Chinese Han population^57^, and its nearby long noncoding RNA, *JAKMIP2-AS1*, promoted the growth of colorectal cancer and indicated poor prognosis^58^. Accordingly, mucin degradation has been linked to intestinal inflammation and tumor incidence^59^. The second strongest signal lied in the gene *SLC9A3* (encoding NHE3: sodium-hydrogen antiporter/exchanger 3; harboring seven rare SVs) associated with the abundance of MF0021: xylose degradation (*p*=1.28×10^−9^). This gene had altered expression in the colonic mucosa of adult IBD patients^60^, showed strong epigenetic interaction with microbiome^61^, and *SLC9A3* knockout mice developed colitis^62^.

In addition to these pathways, some genes were associated with specific microbial taxa. For example, several genes, including *INTS9*, *HLA-DQA1*, and *MTMR3*, were associated with the species *C. intestinalis* from the genus *Collinsella*, which was causally linked to serum cholesterol level^63^ and liver-related disease^64^. The potassium voltage-gated channel subfamily B member 1 (*KCNB1*) SV burden was correlated with the abundance of *Streptococcus salivarius* (p=3.88×10^−7^), *S. vestibularis* (*p*=6.06×10^−7^) and *S. sp.C150* (p=4.23×10^−6^) (**Supplementary Table 7**). A mouse study demonstrated that *S. salivarius* modulates sucrose metabolism via exopolysaccharide (EPS)-dependent short-chain fatty acid production^65^, while *KCNB1* variants impaired pancreatic β-cell function and increased the risk of type 2 diabetes (T2D)^66^. This dual regulation—microbial EPS-SCFA axis and host ion channel activity—suggests a synergistic mechanism whereby *KCNB1* SVs exacerbate T2D risk by disrupting microbial carbohydrate processing and pancreatic glucose sensing. These “host genes-gut microbiome” links need to be further investigated in the future.

### Associations of SVs with blood metabolites

Next, we conducted a GWAS for 22,519 common SVs and 121 blood metabolic traits. A total of 48 associations involving 33 SVs (DEL = 11, INS = 14, DUP = 8) reached genome-wide significance (*p* < 2.2×10^−6^ = 0.05/22519; **Fig. 4a**; **Supplementary Table 8**). Among these, 17 associations reached study-wide significance (*p* < 1.835×10^−8^, adjusting for 22,519 SVs and 121 blood metabolite traits). The genomic inflation factor λ was calculated, with a median value of 1 (ranging from 0.80 to 1.33; **Supplementary Fig. 9**).

**Figure 4.**
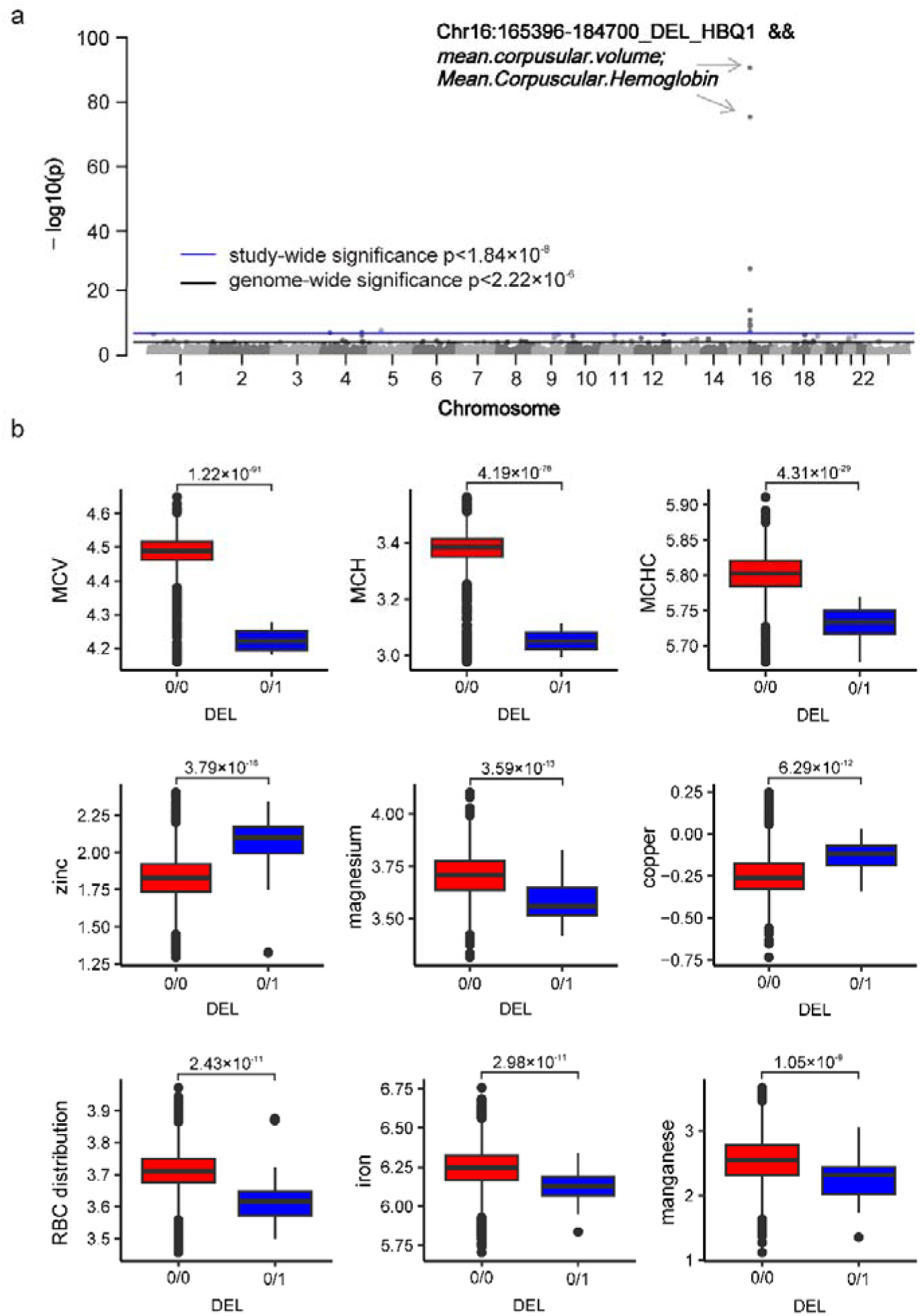
Genome-wide associations of SVs with blood metabolites. (a) Manhattan plot depicting the GWAS results (121 blood metabolite traits; 22,519 common high-quality SVs with MAF ≥ 1%). The study-wide significance (*p* < 1.84 × 10^−8^) and genome-wide significance (*p* < 2.2 × 10^−6^) are indicated by horizontal lines colored blue and black, respectively. The annotations show the DEL variant on chromosome 16 (chr16: 165396-184700), corresponding to the *HBA1*, *HBA2*, *HBM*, and *HBQ1* genes. (b) Boxplots of nine blood metabolite levels associated with the *HBA1/HBA2/HBM/HBQ1* DEL that passed study-wide significance, comparing the metabolite levels between DEL carriers (“0/1”: individuals with DEL) and non-carriers (“0/0”: individuals without DEL). MCV, mean corpuscular volume; MCH, mean corpuscular hemoglobin; MCHC, mean corpuscular hemoglobin Concentration; RBC distribution, red blood cell distribution.

The strongest association signal was observed for a deletion on chromosome 16 (chr16:165396-184700; SV length: 19,304 bp) spanning the *HBA1*, *HBA2*, *HBM*, and *HBQ1* genes, known as the widely reported Southeast Asian deletion (--^SEA^), which is predominantly distributed and associated with anemia in southern China and Southeast Asia^67,68^. In the Chinese population in this study, the MAF of the SEA deletion was 0.01. Consistently, the deletion was mainly detected in East Asian (ASN, MAF=0.019) and South Asian (SAN, MAF=0.013) populations in the 1000 Genomes dataset. The deletion showed remarkably significant associations with nine blood metabolic traits, particularly with mean corpuscular volume (MCV, β = –0.570, *p* = 1.22×10^−91^), mean corpuscular hemoglobin (MCH, β = –0.522, *p* = 4.19×10^−76^), mean corpuscular hemoglobin concentration (MCHC, β = –0.319, *p* = 4.31×10^−29^), and red blood cell distribution (RBC, β = –0.198, *p* = 2.43×10^−11^; **Fig. 4**; **Supplementary Table 8**). Beyond its established correlations with blood cell parameters, the deletion was also associated with blood microelement levels. Carriers of the deletion exhibited elevated zinc and copper concentrations, whereas magnesium, iron, and manganese levels were decreased compared to non-carriers (**Fig. 4b**). In addition to the 16:165396-184700 deletion, a 114 bp duplication on chromosome 9 (chr9:97255555-97255669) within the *SUGT1P4*-*STRA6LP*-*CCDC180* gene was also associated with MCV (β= –0.174, *p* = 7.89×10^−9^) and MCH (β= –0.174, *p* = 7.89×10^−9^) levels. This DUP had an MAF of 0.0125 in this study, which has not been previously reported.

Among the total 48 genome-wide associations, cystine, previously reported as the most heritable metabolite in Chinese^69^, exhibited the highest number of associated SVs, with 16 SVs linked to it (**Fig. 5a**). Three SVs reached study-wide significance, including a 9.2kb deletion on chromosome 5 (5:46048650-46057870; β= –0.163, *p* = 6.47×10^−10^) and two insertions on chromosome 4 (β= –0.161, *p* =2.83×10^−9^) and 12 (β= –0.161, *p* = 2.83×10^−9^), respectively (**Supplementary Table 8**).

**Figure 5.**
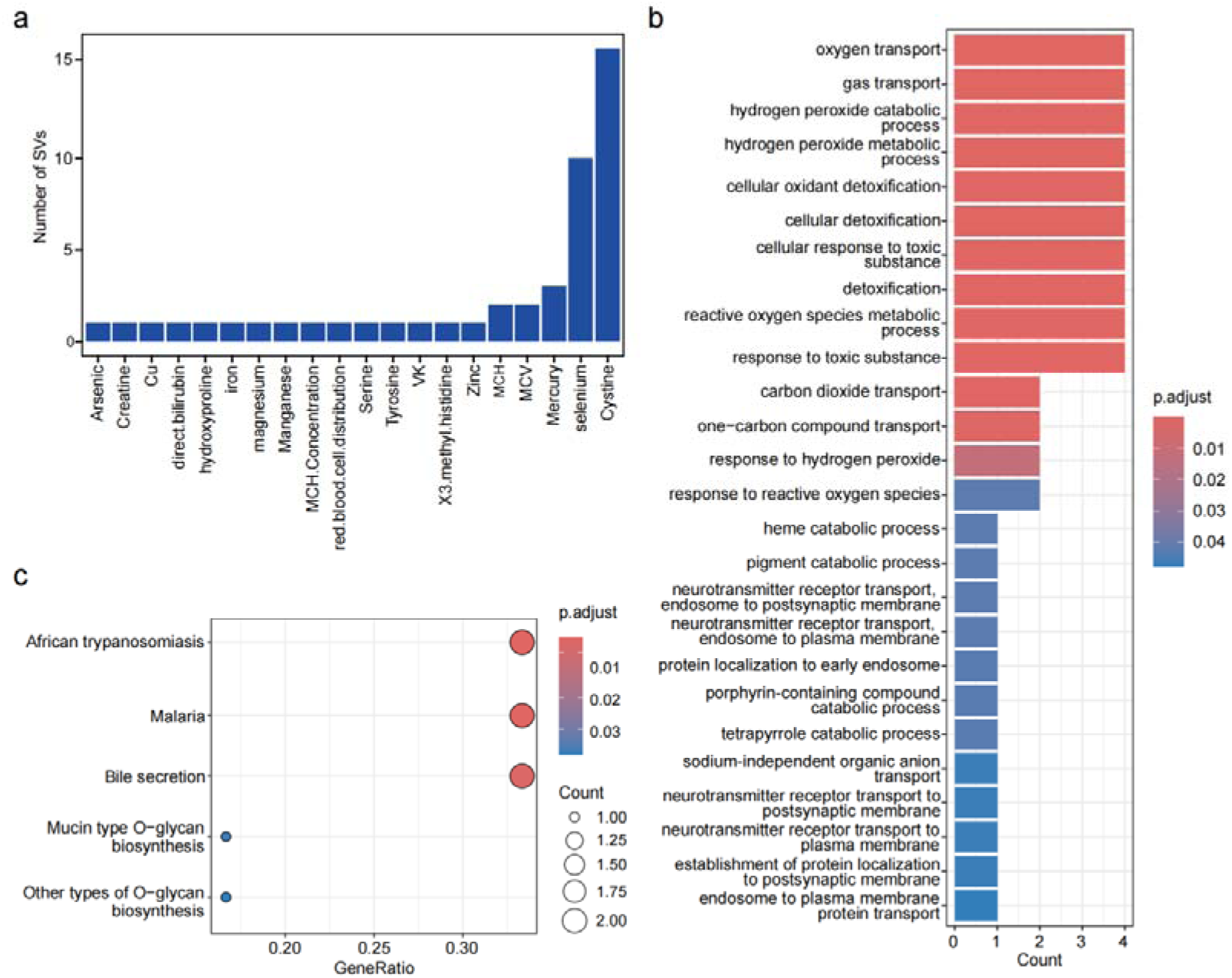
The enrichment analysis of significantly associated SVs in blood metabolites. **(a)** The distribution shows the number of SVs significantly associated with blood metabolites (N=48). (**b and c**) GO (b) and KEGG enrichment analysis (c) for genes harboring SVs significantly associated with the blood metabolites. The bar plot (b) only displays the biological process (BP) results from the GO analysis terms.

The results of GWAS and LD analysis revealed that three out of the 33 SVs appeared to have weaker associations with their respective metabolites compared to neighboring SNPs and could be tagged by neighboring SNPs with r^2^ >0.5 (**Supplementary Fig. 10**). Conditional analysis further confirmed that the associations of the three SVs with metabolites were attenuated and even not significant after adjusting for neighboring SNPs. For example, the *HBA1*/*HBA2*/*HBM*/*HBQ1* deletion was in moderate LD with the lead SNP chr16:199621 across the nine metabolic traits (r^2^=0.76). After conditioning on the SNP, the association of the deletion with these traits was notably weakened, although it remained statistically significant for seven of them (*p*<0.01; **Supplementary Table 8**), highlighting the significance of this SV in influencing host metabolic processes that are not completely accounted for by nearby SNPs. Furthermore, a 64bp insertion (chr19:15866773-15866783) associated with vitamin K concentration was in strong LD with its tagging SNP (r^2^=0.94), while a ∼6kb insertion (chr12:20861159-20861190) associated with the direct bilirubin level was in moderate LD with its tagging SNP (r^2^=0.76). The associations between the two insertions and metabolites were not significant after adjusting for neighboring SNPs. Apart from these three SVs, the remaining 30 SVs continued to exhibit significant links with their respective metabolites after conditional analysis, underscoring the potential driven and regulatory roles of these SVs in blood metabolic traits (**Supplementary Table 8-9**).

Gene-based analysis of the rare SVs identified 22 unique genes associated with blood metabolic metabolites and further confirmed the most correlations with cystine (**Supplementary Table 10**). SV burdens involving six genes, including *CLN3*, *ELF2*, *WDFY4*, *RBPMS*, *IL6ST*, and *SORCS3*, were linked to cystine levels. The most significant and the only one passing multiple test correction (p< 7.6×10^−8^ by adjusting for 5861 genes and 121 metabolites) was for *the CLN3* gene, which encodes a protein that is involved in lysosomal function and is prenylated most likely at cysteine 435^70^. The other candidate associations were listed in the **Supplementary Table 10**.

GO enrichment analysis of the gene sets harboring significant SVs identified 26 BP, 16 CC, and 30 MF terms (**Fig. 5b; Supplementary Table 11**). The enriched gene sets predominantly featured biological process pathways encompassing gas transport, detoxification and redox, protein and receptor transport, metabolism, and degradation, indicating that SVs associated with blood metabolites play a pivotal role in these biological processes. Additionally, KEGG enrichment analysis revealed a notable enrichment of the gene sets in pathways associated with infectious diseases (e.g., African trypanosomiasis, malaria), bile secretion, and glycosylation-related metabolic pathways (**Fig. 5c; Supplementary Table 12**), implying that SVs located near these genes played crucial roles in these metabolic pathways.

### Replication of previously reported metabolites-associated SVs

In the present study, we investigated previously reported metabolites-associated SVs to determine their replication. Out of a total of 58 autosomal SVs identified in previous studies^26–28^, 33 were detected in our samples, with 16 showing associations with metabolites (*p*< 0.05; **Supplementary Table 13-15**). Notably, following a stringent Bonferroni correction threshold of 0.00152 (0.05/33), three SVs remained statistically significant. The most robust signal was observed for *HBA1*/*HBA2*/*HBM*/*HBQ1* (16:165396-184701) SV, which was associated with MCV/MCH, as confirmed by our study, the NHLBI TOPMed program, and the study by Wu et al. Additionally, the SV within the growth hormone receptor (GHR) gene, previously linked to the small round epithelial cell (SRC) count in the study by Wu et al., was found to be associated with multiple metabolites, including cystine, aspartic acid, and glutamic acid in our study. Moreover, the duplication involving the T cell receptor alpha variable (TRAV) genes, known to be associated with C-reactive protein (CRP) levels in the study by Chen et al., was also correlated with valine levels in our study. These replicated associations across different studies underscore the importance of some specific SVs in human metabolism, emphasizing the need for further validation in larger populations to confirm additional potential associations and to advance our understanding in this domain.

### Host SVs associated with both microbiome and metabolites help understand their interactions

Furthermore, we investigated the potential role of host SVs to intricate interactions between the gut microbiome and blood metabolites. Initially, we observed that host SVs showed pleiotropic associations with both microbial features and plasma metabolites, demonstrating significant p-values reaching genome-wide significance (*p* < 2.22 × 10^−6^) for one phenotype and *p* < 0.05 for the other. Specifically, out of the 30 SVs significantly linked to the gut microbiome, 29 were found to be correlated with at least one blood metabolite (**Fig. 6a**; **Supplementary Table 16**). Similarly, among the 33 SVs significantly associated with blood metabolites, 31 were also correlated with at least one gut microbial feature (**Fig. 6b**; **Supplementary Table 17**).

**Figure 6.**
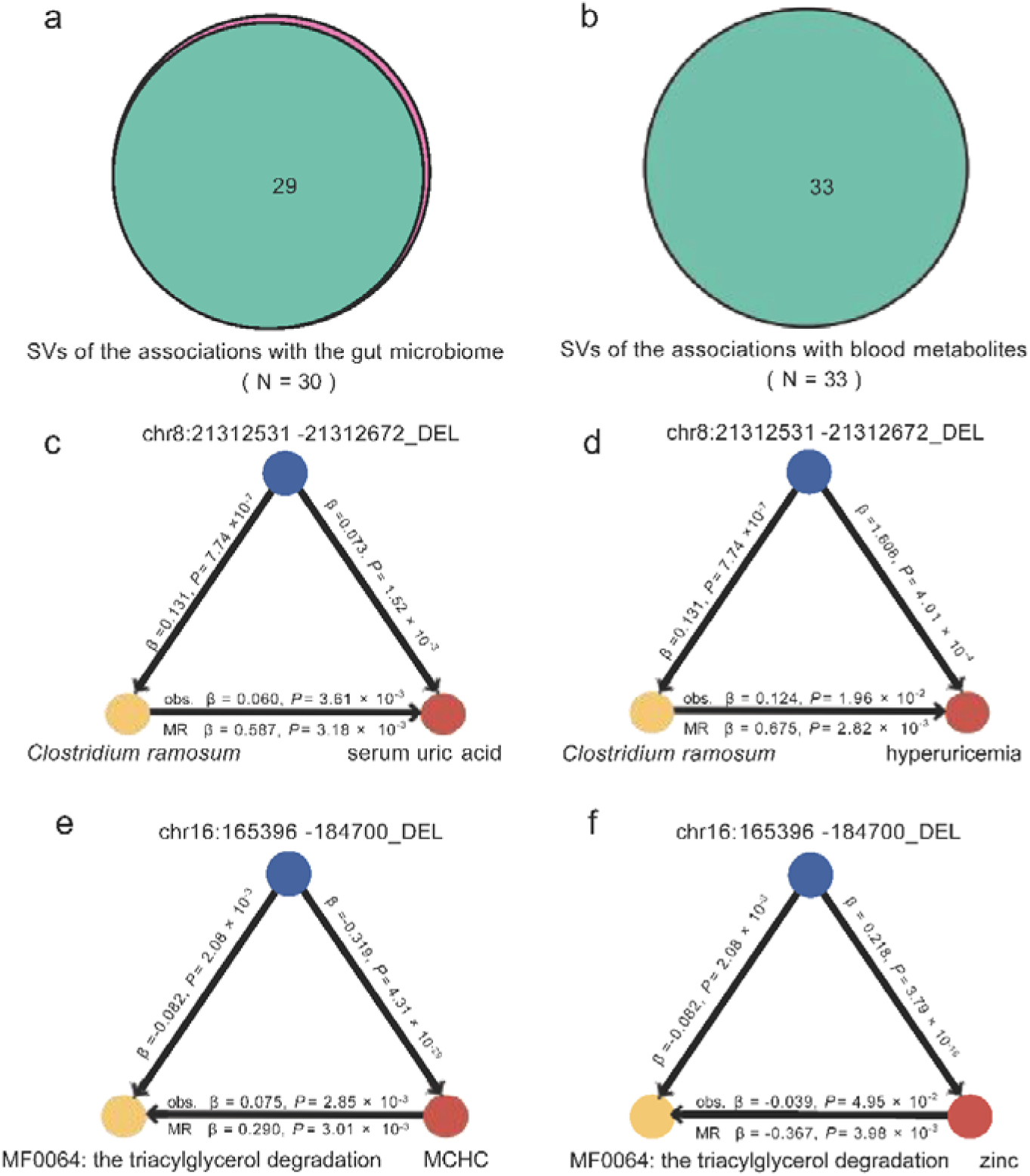
Causal relationships between the gut microbiome and blood metabolites supported by SVs. **(a)** Venn diagrams showing the number of overlapped SVs significantly associated with the gut microbiome (p < 2.2 × 10^−6^) that also linked to blood metabolites (*p* < 0.05). **(b)** Venn diagrams showing the number of overlapped SVs significantly associated with the blood metabolites (p < 2.2 × 10^−6^) that also linked to the gut microbiome (*p* < 0.05). **(c)** The deletion on chromosome 8 (SV length: 141 bp) was significantly associated with both *Clostridium ramosum* (β = 0.131, *p* = 7.47×10^−7^) and the blood metabolite serum uric acid (β = 0.073, *p* = 1.52×10^−3^). A significant observational correlation and further causal effect of *Clostridium ramosum* on serum uric acid supported by MR. **(d)** A significant observational correlation and further causal effect of *Clostridium ramosum* on hyperuricemia supported by MR. **(e)** The causal effect of MCHC on MF0064: the triacylglycerol degradation supported by their overlapped genetic SV, such as the HBA1/HBA2/HBM/HBQ1 deletion (chr16:165396-184700_DEL). **(f)** The causal effect of zinc on MF0064: the triacylglycerol degradation supported by their overlapped genetic SV, such as the HBA1/HBA2/HBM/HBQ1 deletion (chr16:165396-184700_DEL). Obs., observational correlation; MR, Mendelian randomization. MCHC, mean corpuscular hemoglobin Concentration;

Next, we directly calculated the phenotypic correlations between gut microbial features and blood metabolic traits. This approach resulted in the identification of 21 “microbial feature – metabolic trait” pairs meeting stringent significance criteria (associated SV with *p*<2.22 × 10^−6^ for microbiome, *p*<0.05 for the metabolic trait, and *p*<0.05 for correlation between them; **Supplementary Table 16**). For example, the deletion on chromosome 8 (SV length: 141 bp) significantly associated with *Clostridium ramosum* (β = 0.131, *p* = 7.47×10^−7^) was correlated with the level of serum uric acid (β = 0.073, *p* = 1.52×10^−3^). The SV was also linked to an increased risk of hyperuricemia (β = 1.608, *p* =4.01×10^−4^; **Supplementary Table 18**).

Moreover, significant positive correlations were observed between *Clostridium ramosum* and serum uric acid (β = 0.060, *p* = 3.61×10^−3^; **Fig. 6c**) as well as hyperuricemia (β = 0.124, *p*=0.02; **Fig. 6d**). Another instance highlighted an insertion on chromosome 5 (SV length: 96 bp) significantly linked to both *Citrobacter* (β = –0.129, *p* = 1.33×10^−6^) and 1-methylhistidine (β=-0.076, *p* = 4.22×10^−3^), unveiling intricate relationships within *Citrobacter* and 1-methylhistidine (β = 0.048, *p* = 0.023).

Likewise, we also identified 325 “metabolic trait—microbial feature” pairs that met the significance criteria (associated SV with *p*<2.22 × 10^−6^ for metabolic trait, *p*<0.05 for the microbial feature, and *p*<0.05 for correlation between them; **Supplementary Table 17**). For example, the *HBA1*/*HBA2*/*HBM*/*HBQ1* deletion (chr16:165396-184700_DEL), significantly associated with nine blood metabolites such as MCHC and zinc, was also linked to the relative abundance of the microbial pathway MF0064: the triacylglycerol degradation (β = –0.082, *p* = 6.47×10^−10^). A significant association was observed between MCHC and MF0064: the triacylglycerol degradation (β = 0.075, *p* = 2.85×10^−3^; **Fig. 6e**). A significant correlation between zinc and MF0064: the triacylglycerol degradation was also observed (β = –0.039, *p* = 4.95×10^−2^; **Fig. 6f**). These results indicated a potential regulatory network involving microbial and metabolic components, underscoring the importance of considering SVs in the context of microbial/metabolic associations.

To infer causal relationships, we utilized SVs as instrumental variables (IVs) in a Mendelian randomization (MR) analysis. This approach revealed 24 significant causal relationships involving 21 microbial traits and 14 metabolic traits **(Supplementary Table 19**). Observational correlation and MR analysis consistently confirmed the causal effect of *Clostridium ramosum* on increased serum uric acid level and increased risk of hyperuricemia (**Fig. 6c-d)**, as well as that of *Citrobacter* on increased 1-methylhistidine level. Moreover, blood MCHC, zinc, and manganese showed positive causal effects on the microbial pathway MF0064: the triacylglycerol degradation **(Fig. 6e-f)**. Additionally, tyrosine was found to be causally correlated with abundances of diverse gut microbiota, including *Salmonella enterica*, *Bacteroides intestinalis*, MF0052: arginine degradation II, Gammaproteobacteria, *Providencia rettgeri*, MF0093: propionate production I, and Enterobacteriaceae. These results highlight how host SVs could genetically determine the levels of blood metabolites and the gut microbiome, even when the effect of one phenotype was mediated by the abundance of the other phenotype, facilitating a deeper understanding of the interactive mechanisms shaping these biological phenotypes.

## Discussion

This study represents a comprehensive investigation into the role of SVs in the gut microbiome, blood metabolites, and their interactions. Through high-depth whole-genome sequencing of 2,002 individuals, we identified 138,859 high-confidence SVs, nearly half of which were novel. Association analysis revealed 30 significant links between SVs and microbial taxa/pathways, as well as 48 significant associations between SVs and blood metabolites. Among these, 18 reached the study-wide significance. These findings highlight the intricate interaction between host genetic variation and the gut microbiome and metabolites, emphasizing the critical role of SVs in this process. This comprehensive SV dataset serves as a valuable genomic resource for the Chinese population and can function as a reference panel for SVs imputation in GWAS, offering broad utility for future phenotype association studies, including those focusing on diseases.

This study revealed a strong association between specific SVs, particularly a deletion in the *TENT2* gene, and the relative abundance of *Coprobacillus sp. 29_1*. The *TENT2* gene is required for the mono-adenylation of certain miRNAs, such as the liver-specific miRNA-122, which is involved in the regulation of fatty acid homeostasis^71,72^. *Coprobacillus* has been reported to be less abundant and could be a protective species against psoriasis and psoriatic arthritis disease^73–75^. Previous studies have shown that individuals with high *AMY1* copy numbers (*AMY1-CN*) may promote the enrichment of *Ruminococcaceae* microbes (including *Ruminococcus* and *Oscillospira*) by consuming a diet rich in resistant starch, facilitating the efficient fermentation of resistant starch and positively impacting host health^25^. Our analysis identified nine SVs in the *AMY (AMY1, AMY2A, and AMY2B)* gene regions, with one spanning *AMY2A* (DUP, chr1:103611001-103620001), which was significantly associated with *Ruminococcaceae* (β = –0.054, *p* = 0.043). Additionally, a deletion near the *AMY* gene region (DEL, chr1:102630524-102630828) was identified as significantly negatively correlated with *Ruminococcus champanellensis* (β = –0.111, *p* = 1.74 × 10^−5^) and *unclassified Ruminococcaceae bacterium_D16* (β = –0.097, *p* = 2.04 × 10^−4^). Moreover, gene-based analysis further confirmed that *AMY2A/AMY2B* harboring four rare SVs was associated with the microbial pathway of MF0008:maltose degradation. These results consistently supported the correlation between the SV of *AMY* gene region and microbial taxa and pathways, highlighting the role of microbiota in the human genetic adaptation^76^.

Among the “SV-blood metabolite” associations, the most significant association was observed for the 19kb deletion in the *HBA1/HBA2/HBM/HBQ1* gene region. The deletion as the well-known SEA variant exhibited a “thalassemia-like” pattern of red blood cell phenotypic associations, characterized by lower MCH, MCV and RBC. These findings were consistent with previous reports that this deletion is a key SV leading to alpha-thalassemia and is linked to blood cell traits^27,28,77^. In addition to confirming associations with previously reported phenotypes (MCV, MCH, MCHC, and RBC), our findings revealed novel correlations with trace elements, such as copper, zinc, and iron, underscoring the potential utility of this region in disease screening and diagnosis. Beyond the *HBA1/HBA2/HBM/HBQ1* gene region, other associations were partially validated. One study reported that recurrent exon deletions in the haptoglobin (*HP*) gene contribute to lower cholesterol levels^78^. We identified a deletion in the *HP* gene (DEL, chr16:72049453-72076286) significantly associated with total cholesterol (TC, β = 0.061, *p* = 0.04595). We also found an insertion in the *TOX2* gene (INS, chr20:44061952-44061987) significantly associated with TC (β = 0.130, *p* = 2.04 × 10^−5^), LDLC (β = 0.118, *p* = 8.44 × 10^−5^), and HDLC (β = 0.056, *p* = 0.044). The *TOX2* gene encodes a transcription factor involved in immune regulation^79^. Cholesterol and low-density lipoproteins are strongly linked to systemic inflammation^80,81^. This suggests that this SV may influence systemic inflammation and immune function by regulating blood lipid levels, which requires further analysis to clarify the mechanism of this association.

A noteworthy outcome of our study is that the majority of SVs appeared to exert a dominant influence on microbial abundances and metabolic phenotypes, whereas the effects of nearby SNPs appeared to be relatively limited. Traditional genome-wide association studies (GWAS) have primarily focused on the role of SNPs; however, many diseases such as rare and neurodevelopmental disorders^82–84^ have been shown to be caused by SVs. Our study emphasizes the importance of SVs in the gut microbiome and associated metabolite changes, suggesting that future research should not concentrate exclusively on SNP-centric GWAS. Furthermore, most SV loci were found to affect both blood metabolites and the gut microbiota, revealing the important role of SVs in the interplay between microbiota and blood metabolites. Previous studies have established significant observational correlations between the microbiome and host metabolism^85–87^. Our study highlights the crucial role of host genetics at the SV level within this context.

Despite providing valuable insights into SV-microbiota-metabolite interactions, our study had several limitations. Although our sample size was relatively large, particularly for microbiome SV GWAS, further studies are needed in an independent cohort or broader populations to ensure the generalizability of our findings. Moreover, the functional mechanisms of SVs remain incompletely understood. To gain deeper insights into how SVs influence the microbiome and regulate host metabolism, additional research using mouse models or functional experiments is required. As the number of reliable catalogs of SV loci continues to grow, we will gain deeper insights into the impact of host SVs on the gut microbiome and blood metabolites, and a more nuanced understanding of the relationship between the gut microbiota, blood metabolites, and the host’s physiological state. As more robust genetic variants have been identified, future studies should further explore the causal relationships between microbiota and metabolites, thereby elucidating their potential roles in host health and disease.

In summary, this study established a reference SV dataset for the Chinese population and presented compelling evidence regarding the associations of SV with the gut microbiome and blood metabolites, laying the groundwork for future investigations and enriching our knowledge of human health and diseases.

## Methods

### Study cohort and samples

Samples used in this study were obtained from the 4D-SZ omics-cohort, a cohort that has been described elsewhere^14,19,20,29–31^. In the year 2017, a total of 2002 individuals provided blood samples, and 1539 of these individuals also provided matched fecal samples as part of the annual health check program. The blood samples were subjected to high-depth whole genome sequencing and blood metabolic traits testing, whereas the fecal samples were subjected to shotgun metagenomic sequencing. The collection protocols for blood and stool samples, as well as the sequencing procedures, were consistent with those used in earlier studies. Buffy coats were isolated from blood samples, and DNA was extracted using the HiPure Blood DNA Mini Kit (Magen, catalog no. D3111) according to the manufacturer’s protocol. Fecal samples were collected using the MGIEasy kit, and stool DNA was extracted using the MetaHIT protocol. Subsequently, DNA concentrations were quantified using Qubit (Invitrogen), and 200 ng of DNA from each sample was used for library preparation. Paired-end 100bp and single-end 100bp sequencing were performed for blood and stool samples, respectively, on the BGISEQ-500 platform^88^.

Ethical approval for the study was obtained from the Institutional Review Boards (IRB) of BGI-Shenzhen, and all participants provided written informed consent.

### High-depth whole genome sequencing and SNP/indel detection

A total of 2,002 individuals in this study were sequenced to a mean sequencing depth of 42× for the whole genome. The reads were aligned to the human reference genome GRCh38/hg38 using BWA (v.0.7.15)^89^ with the default parameters. The adapter sequences and bases with quality value below five were removed. The alignments were indexed in the BAM format using Samtools (v.0.1.18)^90^ and PCR duplicates were marked for downstream filtering using Picardtools (v.1.62). The Genome Analysis Toolkit’s (GATK v.4.3^91^) BaseRecalibrator created recalibration tables to screen known SNPs and insertions or deletions (INDELs) in the BAM files from dbSNP (v.150). GATKlite (v.2.2.15) was used for the subsequent base quality recalibration and removal of read pairs with improperly aligned segments as determined by Stampy. GATK’s HaplotypeCaller was used for the variant discovery. GVCFs containing SNVs and INDELs from GATK HaplotypeCaller were combined (CombineGVCFs), genotyped (GenotypeGVCFs), variant score recalibrated (VariantRecalibrator) and filtered (ApplyRecalibration). During the GATK VariantRecalibrator process, we took our variants as inputs and used four standard SNP sets to train the model: (1) HapMap3.3 SNPs; (2) dbSNP build 150 SNPs; (3) 1000 Genomes Project SNPs from Omni 2.5 chip and (4) 1000LG phase1 high confidence SNPs. The sensitivity thresholds of 99.9% to SNPs and 99% to INDELs were applied for variant selection after optimizing for Transition to Transversion (TiTv) ratios using the GATK ApplyRecalibration command.

A stringent inclusion threshold for SNP/INDEL variants was established: (i) mean depth >8×; (ii) Hardy-Weinberg equilibrium (HWE) *P* > 10^−5^; and (iii) genotype calling rate > 98%. Furthermore, the sample criteria were as follows: (i) mean sequencing depth > 20×; (ii) variant calling rate > 98%; (iii) no population stratification by performing principal components analysis (PCA) analysis implemented in PLINK (v1.9) ^92^ and (iv) excluding related individuals by calculating pairwise identity by descent (IBD, Pi-hat threshold of 0.1875) in PLINK. Only ten samples were removed after quality control filtering. After variant and sample quality control, 1,992 individuals with 8.5 million common (MAF ≥ 1%) SNP/indel variants were left for association analyses.

### SV detection

Due to the complexity of SV detection from short-read sequencing data, the use of multiple SV calling tools is a common practice to promote accuracy and sensitivity. For SV detection, a pipeline was designed according to the SV-Adjudicator^33^ (https://github.com/talkowski-lab/SV-Adjudicator) and GATK-SV cohort mode (https://github.com/broadinstitute/gatk-sv). In the pipeline, the BAM file is obtained from the preceding alignment and filtering process as its primary input. Subsequently, Manta (v1.6.0)^93^, Wham (v1.7.0)^94^, Svaba (v1.1.3)^95^, and Delly (v0.8.7)^96^ were run for SV detection, xTea^97^ was run for mobile element insertion discovery, and cn.MOPS (v1.40.0)^98^ and GATK gCNV^99^ were run for CNVs calling (**Supplementary Fig. 3**). The algorithms were selected according to Kosugi et al^100^. Finally, the SV calls from the five algorithms and the CNV calls of each sample were standardized and integrated using the GATK-SV pipeline. All computations were completed locally.

### Sample batching and quality control for SV

Following the batching strategy delineated by Collins et al^32^, all samples that passed the quality control (QC) were subdivided into five batches, with ∼380 samples in each batch according to the dosage score (WGD). The distribution of female and male samples in each batch was maintained at a ratio of approximately 1:1, ensuring balance across all batches.

For filtering after GATK-SV integration, outlier samples exhibiting either fewer or higher-than-average SV calls per contig for each caller, or samples manifesting autosomal aneuploidies based on the per-batch binned coverage of each chromosome, were excluded. Additionally, individuals with missing data or those who failed the preceding SNP/indel filtering were omitted. Finally, 1,839 samples were remained in the final SV callset.

### SV filtering

To curate a high-quality SV dataset for subsequent genetic analyses, further filtration of the SVs was conducted post-GATK-SV pipeline execution and sample quality assessment. SVs with quality scores (GQ) > 250 or marked as “PASS” were retained, while variants of type “BND” were excluded. SVs with a genotyping rate >95% and Hardy-Weinberg equilibrium p value > 1×10^−6^ were kept. For GWAS analysis, only SVs with MAF >1% were retained, resulting in a final set of 22,519 SVs left for association analysis.

### SV Annotation

The SVs that passed the quality control were annotated using GATK SVAnnotate (GATK v4.6.0) for functional prediction. The protein-coding GTF file was downloaded from GENCODE (hg38.v37). Only the protein-coding genes were retained for annotation. The BED files of the non-coding regions were downloaded from GATK-SV. AnnotSV (v3.3.8) was applied for additional SV annotation, utilizing the GRCh38 database build to incorporate information such as the pLI score of disrupted genes.

### Validation of SV using NA12878

To validate the SV detection process, the NA12878 human genome reference standard sequenced on the BGISEQ-500 platform (https://db.cngb.org/search/experiment/CNX0094812/) was added as a demo into the cohort for GATK-SV integration. To evaluate the SV detection, the final SV set of the demo was compared with the reference SV sets of NA12878 constructed with the datasets from the Database of Genomic Variants (http://dgv.tcag.ca/dgv/app/home) and the long read-derived SV data^35^. The evaluation was performed using EvalSVcallers (https://github.com/stat-lab/EvalSVcallers).

### Comparisons with published SV datasets

We examined whether the SVs detected in this study have been reported in five publicly available SV databases, including gnomAD, Audano et al., 2019, HGSVC, Wu et al., and dbVar (spanning from 2018 to September 30, 2023). The gnomAD project is a high-coverage (average depth of 32X) short-read sequencing (SRS) dataset comprising 14,891 global samples. The dbVar database catalogs SVs from multiple platforms. We first ensured that the coordinates of the SVs in these public databases were in GRCh38. Our strategy involved the utilization of the R package (v2.3.0) and a custom program to compare our identified SVs with those in public databases, considering a reciprocal overlap rate of 50% or higher.

### Shotgun metagenomic sequencing and profiling

For fecal samples obtained from the SZ-4D cohort, DNA libraries were constructed and sequenced using the BGISEQ-500-DNA sequencer for subsequent metagenomic analysis. Raw data were subjected to quality control and adapter trimming using fastp (v0.23.2) software. To filter out human reads, sequencing reads were aligned to the hg38 reference genome using SOAP2 (v2.22^101^) with an identity threshold of ≥ 0.9. The gene profiles were generated by aligning high-quality sequencing reads to the integrated gene catalog (IGC) by using SOAP2 (identity ≥ 0.95) as previously described^102^. The relative abundance profiles of phylum, order, family, class, genera, and species were determined from the gene abundances. To eliminate the influence of sequencing depth in the comparative analyses, we downsized the unique IGC mapped reads to 20 million for each sample. The relative abundance profiles of genes, phyla, orders, families, classes, genera and species were determined accordingly using the downsized mapped reads per sample.

Gut metabolic modules (GMMs) characterize the bacterial and archaeal metabolism specific to the human gut, emphasizing anaerobic fermentation processes^103^. The current GMM set was developed through a comprehensive review of the literature and metabolic databases, including MetaCyc^104^ and KEGG, followed by expert curation to define modules and alternative pathways. Ultimately, we identified 616 common microbial taxa and GMMs present in at least 10% of the samples.

### Microbiome association analysis with common SVs

We examined the relationship between common host SVs (MAF ≥ 1%) and gut taxa/GMMs by utilizing linear or logistic models based on the abundance of each taxon/GMM. The abundance of taxon/GMM with an occurrence rate exceeding 95% in the cohort was log-transformed, and outliers (deviating over four standard deviations from the mean) were excluded to ensure a reliable quantitative trait analysis. Alternatively, the taxon/GMM was categorized into presence/absence pattern to mitigate zero inflation issues, treating the abundance as a dichotomous trait. Subsequently, for SNP/indel and SV identified in this cohort, we conducted a standard single variant-based mbGWAS analysis using PLINK (v1.9)

^92^ with a linear model for quantitative traits or a logistic model for dichotomous traits. Considering the influence of diet and lifestyle on microbial characteristics, we incorporated age, gender, BMI, defecation frequency, stool form, 12 diet/lifestyle factors, and the top four principal components (PCs) as covariates for mbGWAS analysis.

### Microbiome association analysis with rare SVs

The SV burden test was conducted via generalized linear regression (glm) model with adding age, gender, BMI, defecation frequency, stool form, 12 diet/lifestyle factors, along with the top four PCs as covariates. The gene-based sequence kernel association test (SKAT) analysis was performed in rare SVs (MAF < 0.01) using the SKAT (v2.2.4) package in R with method = ‘skato’. Age, gender, BMI, defecation frequency, stool form, 12 diet/lifestyle factors, and the top four PCs were added into the model as covariates. The rare SVs were filtered using plink v1.9 with options ‘--hwe 1e-4 –-max-maf 0.01 –-geno 0.02 –-mind 0.02’. The gene-based SV annotation was performed as described in SV Annotation section. An SV was annotated within the gene if it overlapped with the gene region or was predicted to affect the promoter of the gene.

### Metabolic trait profiling

In this study, metabolic traits, including anthropometric characteristics and blood metabolites, were assessed in all participants during physical examinations. Anthropometric data, including height, weight, waist circumference, and hip circumference, were recorded by nurses, while age and gender were self-reported. Clinical evaluations, such as blood tests and urinalysis, were conducted by a certified physical examination organization. Hormones were measured in a similar manner, with 250 µl of plasma and atmospheric pressure chemical ionization (APCI). Trace elements were analyzed using a 7700x ICP-MS (Agilent Technologies) equipped with octupole reaction system (ORS) technology to reduce spectral interferences, using 200 µl of whole blood. Water-soluble vitamins were assessed by UPLC coupled to a Xevo TQ-S Triple Quad mass spectrometer (Waters) with ESI in positive ion mode, utilizing 200 µl of plasma. Fat-soluble vitamins were measured with UPLC linked to an AB Sciex Qtrap 4500 mass spectrometer using APCI and 250 µl of plasma. Blood amino acids were quantified using ultra-high pressure liquid chromatography (UHPLC) coupled with an AB Sciex Qtrap 5500 mass spectrometer, utilizing electrospray ionization (ESI) in positive ion mode with 40 µl of plasma. Hyperuricemia is defined as an elevated serum uric acid level, with greater than 6 mg/dL in women and 7 mg/dL in men.

### Metabolites association analysis

The log10-transformed median-normalized values of 121 anthropometric and metabolic traits were used as quantitative traits. Samples with missing values or values greater than 4 standard deviations from the mean were excluded from the association analysis. Each of the common SNP/indel or common SV (MAF ≥ 1%) was independently tested using a linear model for quantitative traits in PLINK v1.9. The gene-based sequence kernel association test (SKAT) analysis was performed for rare SVs (MAF < 0.01) using the SKAT (v2.2.4) package in R, as done in the microbiome association analysis. Covariates included age, gender, and the top four PCs.

### Gene enrichment analysis

The genes harboring significantly associated SVs were annotated using AnnotSV^54–56^. Gene Ontology (GO) and Kyoto Encyclopaedia of Genes and Genomes (KEGG) gene functional enrichment analysis of genes harboring significantly associated SVs were executed by using the R packages including clusterProfiler, enrichplot, ggplot2 and org.Hs.eg.db. A standardized metric (*p*-value < 0.05, Q-value < 0.2) was used to prioritize the top functional items and pathways. Biological process (BP), cellular component (CC), and molecular function (MF) were the three categories included in GO analysis.

### Observational correlation analyses

All microbial features and metabolic traits were transformed using log-transformed median-normalized function to reduce the skewness of the distributions. The observational correlations between these variables were evaluated using multivariable linear regression analysis while adjusting for age and gender. The beta coefficient and p-value were obtained.

### One-sample MR analysis

To explore the potential causal relationships between microbial traits and metabolic characteristics within the same cohort, we conducted a one-sample bidirectional MR analysis in this cohort with both metabolite and microbiome data. We selected SV instruments based on a stringent genome-wide significance threshold (*p* < 2.22 × 10LL) and performed clumping with an LD r² < 0.1 to ensure independence among genetic variants for MR analysis. An unweighted polygenic risk score (PRS) was calculated for each participant based on independent genetic variants derived from GWAS data. For each SV, we coded the risk alleles as 0, 1, or 2, depending on the number of trait-specific risk alleles carried by the individual. Instrumental variable (IV) analyses were conducted using two-stage least squares (TSLS) regression. In the first stage, we assessed the association between the PRS and the observational phenotype using linear regression, from which we derived fitted values representing the predicted exposure. During this process, the *F*-statistics and the variance explained by IVs were calculated. We required the *F*-statistics over 10 to ensure a strong IV for analysis. In the second stage, we regressed the outcome trait on the genetically predicted exposure from the first stage. Both stages were adjusted for potential confounders, including age, gender, and BMI. TSLS was performed using the ‘ivreg’ function from the AER package in R (v.1.2-9; https://www.rdocumentation.org/packages/AER/versions/1.2-9/topics/ivreg).

## Data Availability

All summary statistics that support the findings of this study including the population SV dataset were available in the **Supplementary Tables**. The gut metagenomic sequencing data after removing host reads have been publicly available in the GSA from https://ngdc.cncb.ac.cn/gsa-human/browse/HRA000907. All the metabolites and individual-level host genetics data have been uploaded to the GSA database (https://ngdc.cncb.ac.cn/bioproject/browse/PRJCA005334). According to the Human Genetic Resources Administration of China regulation and the institutional review board of BGI-Shenzhen related to protecting individual privacy, these data are controlled-access and are available via an application on request. The human reference hg38 datasets are publicly available from http://hgdownload.soe.ucsc.edu/goldenPath/hg38/bigZips/. The datasets of common SVs are available at the following link: GnomAD SV: https://datasetgnomad.blob.core.windows.net/dataset/release/4.1/genome_sv/gnomad.v4.1.sv. sites.vcf.gz, dbvar SV: https://gnomad.broadinstitute.org/, HGSVC SV: http://ftp.1000genomes.ebi.ac.uk/vol1/ftp/data_collections/HGSVC2/release/v1.0/integrated_callset/freeze3.sv.alt.vcf.gz, Wu et al SV: https://ftp.ncbi.nlm.nih.gov/pub/dbVar/data/Homo_sapiens/by_study/csv/all_variants_for_nstd212.csv.gz, Audano et al. 2019, http://ftp.1000genomes.ebi.ac.uk/vol1/ftp/data_collections/hgsv_sv_discovery/working/20181025_EEE_SV-Pop_1.

## Author Contributions

X.L and T.Z. conceived and organized this study. X.X., H.Y., Y.Z., W.L, X.J., and L.X. led the organization of the cohort, the sample collection, and the questionnaire collection. M.H. and H.L led the DNA extraction and sequencing. X.L and X.T. processed the whole genome data. L.Z., Y.J., and M.L. processed the metagenome data. X.L., X.T., and L.Z. performed the metagenome-genome-wide association analyses and Mendelian randomization analyses. X.L. wrote the raw manuscript. All authors contributed to data and texts in this manuscript.

## Competing interests

The authors declare no competing interest.

## Acknowledgments

We sincerely thank the support by the National Natural Science Foundation of China (No. 32200548) and the China National GeneBank. We thank all the volunteers for their time and for self-collecting the oral samples using our kit. The schematic workflow diagram of the study was drawn by Figdraw (https://www.figdraw.com).

## Notes

### Competing Interest Statement

The authors have declared no competing interest.

### Summary of Updates

Manuscript revised; figure 6 revised; graphic abstract image revised

